# Evolution of connections and cell types: integrating diffusion MR tractography with gene expression data highlights increased cortico-cortical projections in primates

**DOI:** 10.1101/493593

**Authors:** Christine J. Charvet, Arthi Palani, Priya Kabaria, Emi Takahashi

## Abstract

Diffusion MR tractography permits investigating the three-dimensional structure of cortical pathways as interwoven paths across the entire brain. We use high-resolution scans from diffusion spectrum imaging and high angular resolution diffusion imaging to investigate the evolution of cortical pathways within the euarchontoglire (i.e., primates, rodents) lineage. More specifically, we compare cortical fiber pathways between macaques (*Macaca mulatta*), marmosets (*Callithrix jachus*), and rodents (mice, *Mus musculus*). We integrate these observations with comparative analyses of Neurofilament heavy polypeptide (NEFH) expression across the cortex of mice and primates. We chose these species because their phylogenetic position serves to trace the early evolutionary history of the human brain. Our comparative analysis from diffusion MR tractography and NEFH expression demonstrates that the examined primates deviate from mice in possessing increased long-range cross-cortical projections, many of which course across the anterior to posterior axis of the cortex. Our study shows that integrating gene expression data with diffusion MR data is an effective approach in identifying variation in connectivity patterns between species. The expansion of cortico-cortical pathways and increased anterior to posterior cortical integration can be traced back to an extension of neurogenetic schedules during development in primates.

## Introduction

One of the fundamental issues in evolutionary neuroscience is to understand the principles with which connections evolve (Nudo and Masterdon, 1988, 1990; Barbas H, Rempel-Clower, 1997; Striedter, 2005; Kaas, 2012; Hofman, 2014; Barbas, 2015; Finlay, 2016; Horvát et al., 2016; Hilgetag et al., 2016; Das and Takahashi, 2017; Beul et al., 2017; Goulas et al., 2018). Diffusion MR imaging has begun to answer long-standing questions in the field of evolutionary neurobiology (Rilling et al., 2008; Takahashi et al., 2011; Charvet et al., 2017a; Das et al., 2017; Mortazavi et al., 2018). For instance, diffusion MR tractography has demonstrated that the human arcuate fasciculus is expanded relative to that of other primates, which might have been an important pathway responsible for the emergence of language in humans (Rilling et al., 2008; Rilling, 2014). More recently, the use of diffusion MR tractography has demonstrated that primates possess increased anterior to posterior cortical integration compared with a number of other mammals (Charvet et al., 2017a). We use diffusion MR tractography to investigate the evolutionary history of cortical pathways across euarchontoglires (i.e., primates, rodents) to trace the early evolutionary history of the primate lineage leading to the human brain.

Diffusion MR imaging is a high-throughput and non-invasive method that is instrumental in investigating long-range projection pathways coursing across the white matter of the brain. Questions focused on the evolution of connections have traditionally been addressed with tract-tracers, which are time-consuming, invasive, and unavailable to the study of the human brain (Gilbert et al., 1975; Kennedy and Bullier, 1985; Barbas, 1986; Nudo and Masterton, 1990; Hof et al., 1995). We here integrate observations from diffusion MR tractography with comparative analyses of gene expression data from bulk and single cells to trace the evolution of cortical pathways in euarchontoglires. We focus especially on species differences in the expression of Neurofilament heavy polypeptide (NEFH) because NEFH is preferentially expressed by large neurons with long axons projecting over long distances (Hof et al., 1995; Marszalek et al., 1996; Friedland et al., 2006; Nguyen et al., 2017). Therefore, comparative analyses of NEFH expression levels coupled with observations from diffusion MR tractography can be used to identify evolutionary changes in long-range projection patterns (Hof et al., 1995).

The location of a neuronal soma across the depth of the cortex predicts projection patterns to some extent. Variation in neuron numbers across the depth of the cortex can identify evolutionary changes in connectivity patterns. Although little is known about the extent to which these stereotypical projection patterns vary across species, neurons located superficial to the cortical surface (i.e., upper layer neurons, layers II-II) preferentially project cross-cortically. Layer IV neurons are local circuit neurons. This is in contrast with neurons located towards the white matter (i.e., lower layer neurons, layer V-VI) that preferentially, though not exclusively, project sub-cortically (Gilbert et al., 1975; Kennedy and Bullier, 1985; Barbas, 1986; Nudo and Masterton, 1990). That is, some layer V neurons may form long-range, short-range or collosal pathways (Barbas 1986; Barbas and Rempell-Clower 1997; Barbas et al 2005ab; Hilgetag and Grant, 2010). Primates possess disproportionately more layer II-IV neurons compared with many other mammals, including rodents (Charvet et al., 2015, 2017a,b,c). Although the relative contribution of layer II-III versus layer IV neuron numbers has not been systematically quantified across the cortex of different species, the observation that primate cortices possess an expansion of layer II-IV neuron numbers suggests that primates may possess increased cross-cortically projecting neurons compared with rodents. However, it remains unclear whether the increase in upper layer neuron numbers in primates is concomitant with increased long-ranging cortical projections, local circuit neurons, or both. Tract-tracing studies and diffusion MR studies have reported that primates possess a large number of long-range cross-cortically projecting neurons across the anterior to posterior axes of the cortical white matter (Schmahmann et al., 2007; Schmahmann and Pandya, 2009; Charvet et al., 2017a). Given these reports, the aim of the present study is to test whether primates do indeed possess more long-range cross-cortically projecting neurons compared with other species such as mice.

We integrate comparative analyses of NEFH expression with diffusion MR tractography to test for deviations in long-range cortically projecting neurons in primates and in rodents. We select to compare brain fiber pathways between macaque (*Macaca mulatta*, Pandya and Kuypers, 1969; Schmahmann et al., 2007; Schmahmann and Pandya, 2009), marmoset (*Callithrix jacchus,* Okano and Mitra 2015), and mice (*Mus musculus*, Wu et. al. 2013), in part, because brain pathways have been mapped in earlier studies with the use of tract-tracers and MRI diffusion tractography in these species. We also select these species because of their phylogenetic position. Rodents (e.g., mice) are the closest living-outgroup to primates and marmosets are phylogenetically intermediate between macaques and rodents (Bininda-Emonds et al., 2007; Perelman et al., 2011). Comparative analyses of pathways between rodents and primates serve to trace the early evolutionary history of the human brain.

The main finding from the present analysis is that there are major modifications to cortical pathways between primates and mice. Integrating comparative analyses from diffusion MR tractography and NEFH expression patterns demonstrate that primates possess an expansion in long-range cortico-cortical pathways compared with mice. Many of the identified cortico-cortical long-range projection pathways course across the anterior to posterior axes of the cortex, suggesting that the primate cortex deviates from other species in possessing increased anterior to posterior cross-cortical integration.

## Materials and Methods

### Specimen Preparation

Brains of species used for the present study were collected opportunistically from a group involved in vision research at Harvard Medical School (HMS). All procedures were approved by the Institutional Review Board (IRB) at Massachusetts General Hospital and HMS where our MRI scans were performed. No animals were killed for the purpose of this study. We used brains of four adult macaques, three marmoset monkeys, and four mice. All procedures were approved by the Massachusetts General Hospital and HMS. Briefly, the brains of macaques, marmosets, and mice were perfused with phosphate-buffered saline solution followed by 4% paraformaldehyde, and fixed for 1 week in 4% paraformaldehyde solution containing 1 mM gadolinium (Gd-DTPA) MRI contrast agent to reduce the T1 relaxation time while ensuring that enough T2-weighted signal remained. For MR image acquisition, the brains were placed in a Fomblin solution (Fomblin Profludropolyether; Ausimont). Specimens were scanned with a 4.7 T Bruker Biospec MR system at the A.A. Martinos Center for Biomedical Imaging. The brains of 2 human adults without neurologic disease were from autopsy. The autopsies were performed in with consent of the family and included permission for research under our IRB-approved protocol.

### Scanning Parameters

For all scans, we determined the highest spatial resolution for each specimen with an acceptable signal-to-noise ratio of more than 130. For mice and marmosets, the pulse sequence used for image acquisition was a 3D diffusion-weighted spin-echo echo-planar imaging sequence at a 4.7 T Bruker Biospec MR system with a high-performance gradient and a radio-frequency coil that better fits the small brains. We used TR 400 msec, TE 20 msec, with number of segments 2. Spatial resolution was 0.13 × 0.13 × 0.17 mm for mice, and 0.23 × 0.23 × 0.28 mm for marmoset. Sixty diffusion-weighted measurements (b = 4,000 sec/mm^2^) and one non diffusion-weighted (b = 0) measurement were acquired, with δ= 12.0 msec, Δ = 24.0 msec as previously described (Rosen et al., 2013; Fame et al., 2016; Kanamaru et al., 2017). The total acquisition time was about 2 hours and 10 minutes for each imaging session.

Human brains were scanned on a 3-Tesla TimTrio MRI scanner (Siemens Medical Solutions, Erlangen, Germany) using a 32-channel head coil. Diffusion data were acquired using a 3-dimensional diffusion-weighted steady-state free-precession sequence (McNab et al., 2009) that used 44 diffusion-weighted measurements. Spatial resolution was 1.0 x 1.0 x 1.0 mm. Total scan time for each specimen was 5.5 hours. Additional diffusion sequence parameters have been previously reported (Edlow et al., 2012; Kolasinski et al., 2013).

We performed diffusion spectrum encoding as previously described for macaques (Wedeen et al. 2005). Briefly, we acquired 515 diffusion-weighted measurements, corresponding to a cubic lattice in q-space contained within the interior of a ball of maximum radius bmax = 4 × 10^4^ cm^2^/s, with δ = 12.0ms, Δ = 24.2ms. The total acquisition time was 18.5 hours for each experiment. Spatial resolution was 0.55 × 0.55 × 0.75 mm. The total acquisition time was about 18 hours. Some of these scans were used in a previous study (Charvet et al., 2017a).

### Diffusion Data Analyses—Tractography

We used Diffusion Toolkit to utilize a streamline algorithm for diffusion tractography. Trajectories were propagated by consistently pursuing the orientation vector of least curvature. We terminated tracking when the angle between 2 consecutive orientation vectors was greater than the given threshold (45°) or when the fibers extended outside the brain surface, by using mask images of the brains created by TrackVis (http://trackvis.org) for each specimen. In many tractography studies, FA values are used to terminate fibers in the gray matter, which in adults has lower FA values than the white matter. As one of the objectives of our study was to detect fibers in low FA areas, we used brain mask volumes to terminate tractography fibers without using the FA threshold for tractography, which is an approach consistent with that of previous studies (Takahashi et al., 2010, 2012). Trajectories were displayed on a 3D workstation of TrackVis. The color-coding of fibers is based on a standard RGB code, applied to the vector between the end-points of each fiber (**Figs. 1, 2**).

**Figure 1.**
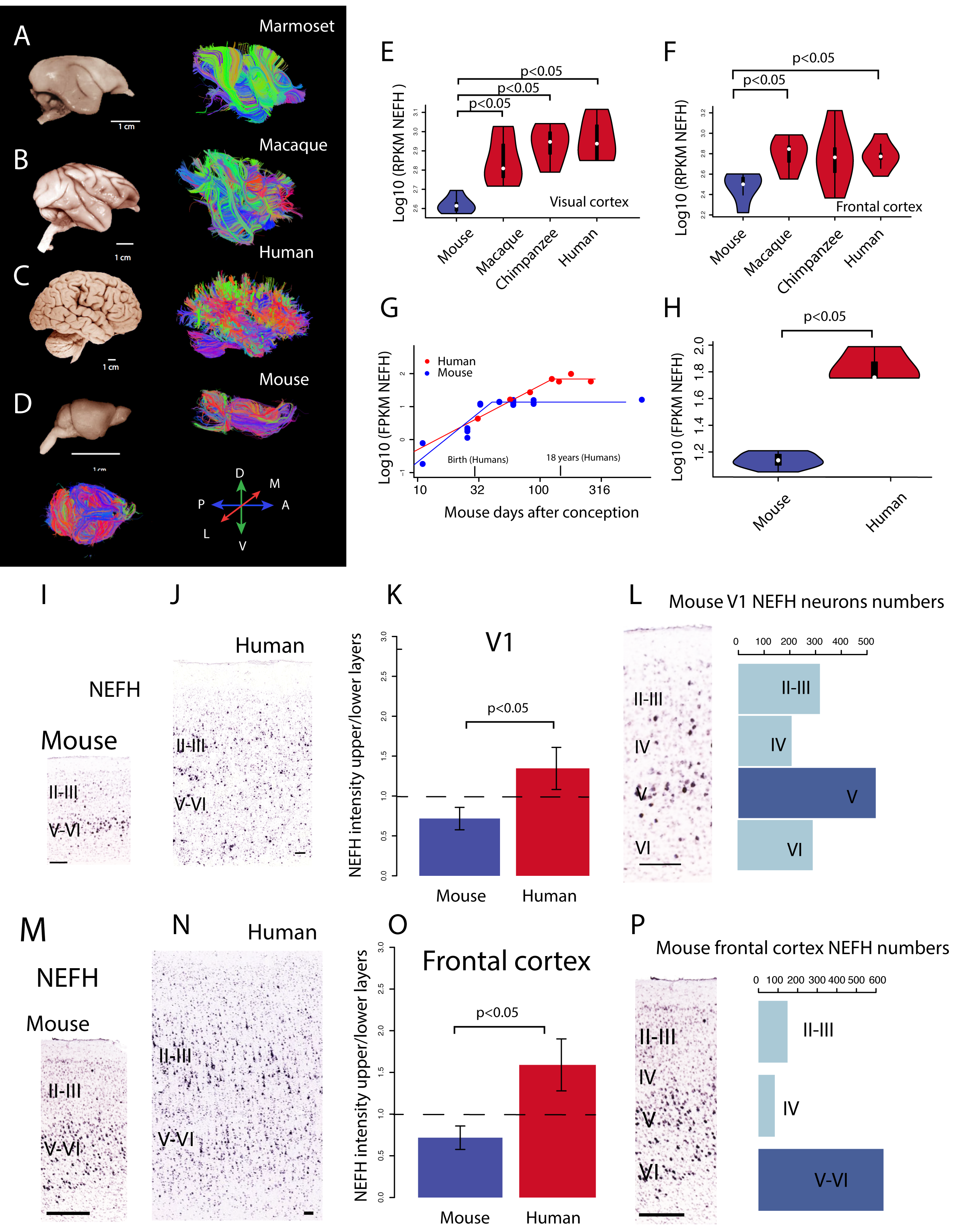
Brain pictures and overall tractography pathways of marmosets (A), macaques (B), humans (C), and mice (D). For tractograpy, axial filters were set through the middle of the brain at the level of the thalami. Tractography of marmosets (A), macaques (B), and humans (C) show a number of inter-cortical pathways coursing across various directions through the cortex. In contrast, mice (D) possess inter-cortical pathways nearly exclusively coursing across the anterior to posterior axis or the medial to lateral axis. We compare NEFH expression values because it is a marker of long-range connecting cortical neurons (Hof et al., 1995). (E-F) A box plot highlighting the median (white dot), quantiles, and confidence intervals overlaid on a violin plot (i.e., density plot) show the distribution of NEFH expression across individuals for each species in the visual cortex (E) and the frontal cortex (F). NEFH expression is significantly higher in the visual cortex (E) of humans, chimpanzees, macaques compared with mice. A similar situation is observed in the frontal cortex (F) where NEFH expression is significantly increased in humans and macaques compared with mice. (G) Developmental time course of NEFH expression shows major differences in NEFH expression between humans and mice. NEFH expression is plotted against equivalent developmental time points in humans and mice on the axis where age of mice is mapped onto humans (Workman et al., 2013). (H) NEFH expression increases and reaches a plateau in humans and in mice but the plateau in NEFH expression is reached much later than expected in humans after controlling for overall changes in developmental schedules. (H) Data from (G) shows that NEFH expression is significantly greater in humans compared with mice once NEFH expression reaches a plateau. (I-J) NEFH mRNA expression is more superficially expressed in humans at 17 years of age compared with mice at post-natal day 56. (K) The density of NEFH found in upper relative to that observed in lower layers in the primary visual cortex of humans (17-57 years old) is significantly greater in humans than in mice (postnatal-day 28 to 24 months old). (L) Single cell RNA sequencing from the mouse primary visual cortex show that there are relatively fewer expressed NEFH labeled cells in upper versus lower layers. These observations are consistent with the notion that NEFH expression is much lower in upper layers relative to lower layers in mice. A similar situation is observed in the frontal cortex (M-P) where NEFH is relatively increased in upper versus lower layers in humans (aged 28 year old) relative to mice (P56; M-O). (P) Single cell RNA sequencing analyses with dropviz (http://dropviz.org/) highlights that most of the NEFH-labeled neurons are located in lower layers. Abbreviations: A: anterior; P: posterior; M: medial; L: lateral, D: dorsal; V: ventral; FPKM: fragments per kilobase million. (I-J) Sections of NEFH mRNA expression are from the Allen brain atlas 2010 Allen Institute for Brain Science. Allen Human Brain Atlas. Available from: human.brain-map.org. Scale bar: 100 micron.

**Figure 2.**
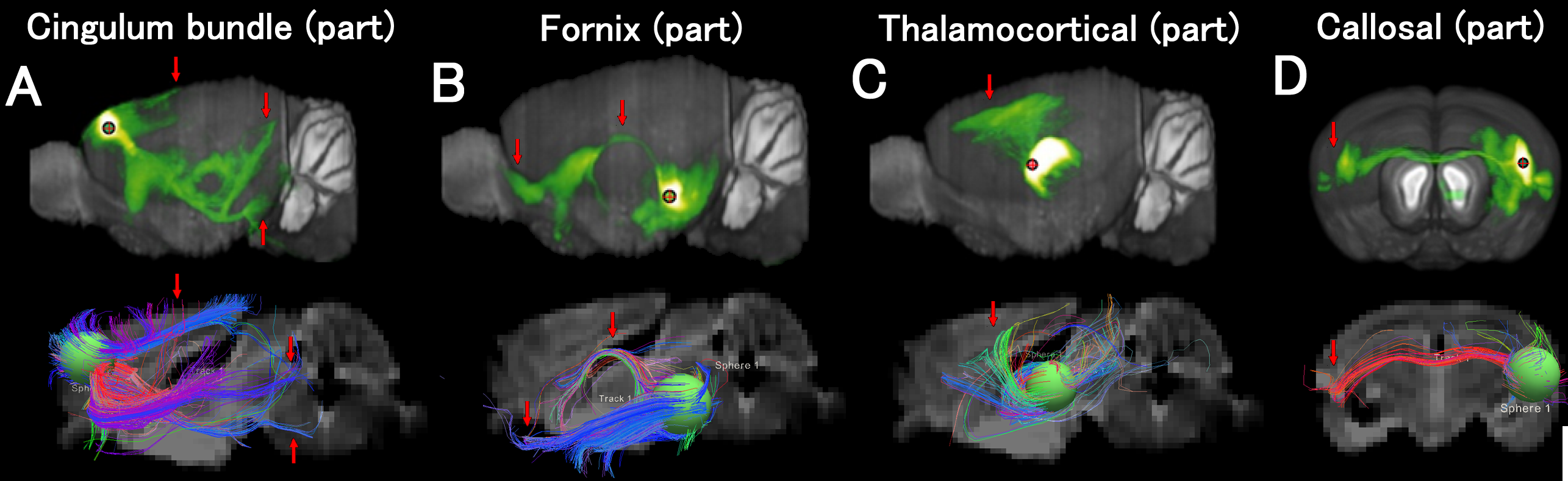
Comparative analyses of tractographies from tract-tracers (upper panels) and diffusion MR scans (lower panels) in mice. Data from tract-tracers are from the Allen Brain Atlas (ABA) mouse brain connectome, which is one of the most comprehensive data-sets to map mouse brain connectivity, from anterograde viral neuronal tracers (Oh et al. 2014). These analyses highlight strong concordance in tractography for the cingulum bund (A), the fornix (B), thalamo-cortical pathways (C), as well as the corpus callosum (D) across the two methods.

### Diffusion Data Analyses—ROI Placement and Restriction Parameters

In Trackvis, we used frames set through the hemisphere with a minimum length threshold of 10-15 mm in primates and 5 mm in mice to highlight cortical pathways (**Fig. 1**), This approach offers a global perspective with which to examine pathways across the brain of different species. We also include a diffusion MR scan of a human brain. Details of procedures for data acquisition have been published previously (Charvet et al., 2017a).

Using ITK-SNAP (http://www.itksnap.org/), we were able to segment the three-dimensional images of each of the brains respectively in order to create a visual representation of the cortical surfaces. We then used TrackVis in order to visualize diffusion tractography pathways and associated diffusion weighted images as an anatomical reference. Regions of interest (ROIs) were created for each gyrus and associated sulcus. In both macaques and marmosets, multiple spherical ROIs were used for the orbitofrontal, superior temporal and temporal gyri and sulci, and hand-drawn ROIs were used for the calcarine gyrus and sulcus due to the complex structure and depth of the sulcus in the macaque. We did not use length thresholds to identify these pathways.

In order to compare the cingulum bundle, fornix, and thalamocortical/corticothalamic pathways across all three species, ROIs were placed following the method previously published (Catani and de Schotten, 2008; Song et al., 2015; Cohen et al., 2016). The selected fibers were further filtered for their length with the software program TrackVis. Minimum length thresholds of 58 mm, 15 mm, and 13 mm were utilized for macaques, marmosets, and mice, respectively. In marmosets, a minimum threshold of 15 mm was used in order to isolate pathways of interest. What may constitute equivalent length thresholds between species may be debatable, considering that the brains of the examined species vary in size and length of pathways.

We confirmed our mouse tractography results with tract tracer studies available in the Allen Brain Atlas (http://portal.brain-map.org/) (**Fig. 2)**. Tract-tracer data are from the Allen Brain Atlas (ABA) mouse brain connectome, which is one of the most comprehensive data-sets to map mouse brain connectivity, from anterograde viral neuronal tracer injections in the wild-type C57BL/6 mouse brain (Oh et al. 2014). We use these data to ensure the condordance of diffusion MR tractograpjy with tractographies from tract-tracers. Injection sites in the Allen Brain Atlas that most closely identify the cingulum bundle (A), fornix (B), thalamocortical (C), and callosal (D) pathways were selected, and ROIs (1mm in radius) for tractography were placed in the injection sites; the dorsal anterior cingulate cortex (**Fig. 2A**) the hippocampal C1 region (**Fig. 2B**), and the reticlular nucleus of the thalamus (**Fig. 2C**), which were corresponding ROI regions for the images from the Allen Brain Atlas (**Fig. 2, upper rows**).

### Comparative analyses of NEFH expression

Our comparative analysis from diffusion MR tractography suggests that the cortex of primates possess alterations in cross-cortically projecting pathways compared with mice (see also Charvet et al., 2017a). To validate these observations quantitatively, we compare the expression of Neurofilament Heavy Polypeptide (NEFH) in primates and mice. NEFH is preferentially expressed by neurons with thick myelinated and long axons with high conduction velocities (Marszaelk et al., 1996; Usoskin et al., 2015). Thus, NEFH can be used to assess species differences in cortical long-range projection patterns (Hof et al., 1995; Friedland et al., Nguyen et al., 2017). We test whether NEFH is differentially expressed in the prefrontal and visual cortices of mice (n=5; post-natal day 56) and primates (macaques n=5; 9-21 years of age; chimpanzees n=6, 21 to 42 years of age; humans, n=5, 21-59 years of age; GEO accession number: GSE49379, Bozek et al., 2014). NEFH expression values are in reads per kilobase million (RPKM). Although NEFH has not been compared quantitatively between primates and rodents, there are qualitative differences in NEFH expression across the depth of the cortex between humans and mice (Zeng et al., 2012). However, Zeng et al., 2012 focused exclusively on the primary visual cortex of humans and mice, and it is not clear whether species differences in NEFH expression are restricted to the primary visual cortex, or whether these differences in NEFH expression span cortical areas other than the primary visual cortex. In primates, NEFH expression appears preferentially expressed in upper layers in some cortical areas but not in others (Barbas and Garcia-Cabezas 2016).

For each species, we performed a principal component analysis on the log-based RPKM values of orthologous and expressed genes across the four species under study in order to detect and remove outliers. The removal of an outlier was based on whether a sample deviated from most other samples on principal component 1 and 2. We removed a total of 5 outliers from all samples from the prefrontal and visual cortices. We used a Kolmogorov–Smirnov test (ks test) to explore differences in NEFH expression between species. We selected a ks test because it does not assume that the data is normally distributed.

To investigate potential species differences in developmental trajectories of NEFH, we use a previously published RNA sequencing dataset from bulk samples of the frontal cortex of humans (mid-frontal gyrus; n=7) from around birth to 55 years of age and mice from embryonic day 11 to 22 months of age (frontal cortex; n=21; Lister et al., 2013). NEFH expression values are in fragments per kilobase million (FPKM; GEO accession number: GSE47966; Lister et al., 2013). We report boxplots overlaid on density plots (i.e., violin plots) to compare the distribution of NEFH expression across species (**Fig. 1**). We use a model that relies on the timing of transformations across mammalian species to find equivalent stages of development between model organisms and humans (Workman et al., 2013). A total of 271 developmental transformations were acquired across 18 mammalian species (e.g., mice, rats, rhesus macaques, humans). Developmental transformations capture rapid changes that occur during brain development. Many of these developmental transformations focus on changes in synaptogenesis, neuron production, as well myelination. The timing of developmental transformations can be used to identify corresponding ages across species. To that end, the timing of development transformations (logged-transformed) are regressed against an event scale in each species. The “event scale” is a multivariate measure of overall maturational state of the nervous system, with the generation of the first neurons near “0,” with “1” corresponding to about 2 years postnatal in humans. We can use these data to identify whether select developmental processes occur unusually early or late in a given species after controlling for overall changes in developmental schedules (Clancy et al., 2001; Dyer et al., 2009; Charvet et al., 2017b, 2018l See **Fig 3, 4, 5**). Because the translating time model extends up to two years of age in humans and its equivalent in mice, we extrapolate corresponding ages at later time points. To that end, we identified equivalent ages from a linear regression of the logged-based 10 values of days after conception for humans and mice. This permitted extrapolating corresponding ages between the two species throughout adulthood in the two species (**Fig. 1**, **3**). We identified when NEFH expression reaches a linear plateau with the library package easynls (model 3) in R.

**Figure 3.**
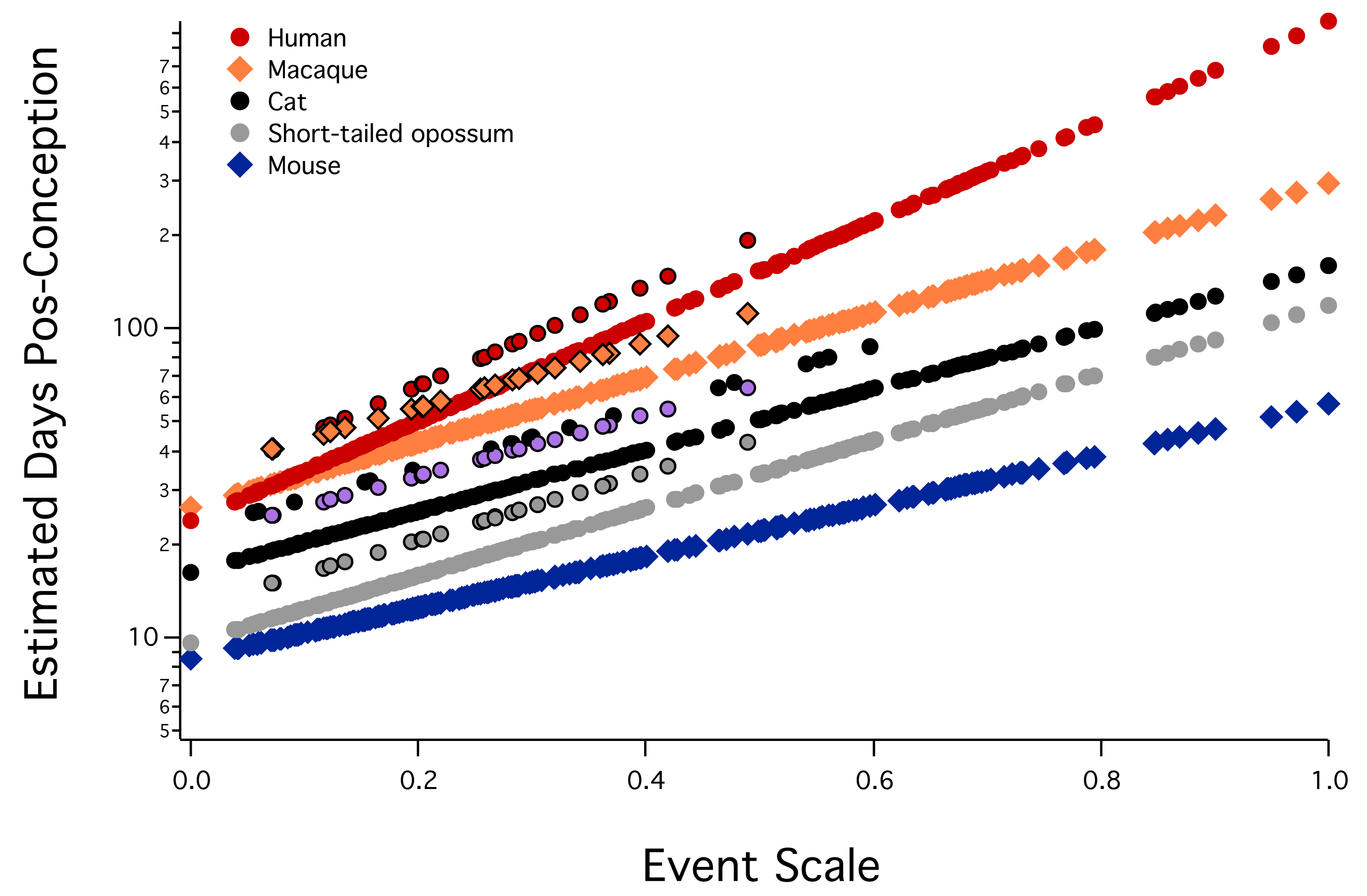
(A) Estimated days after conception (log-transformed) is plotted against an event scale in some of the 18 mammalian species previously studied (i.e., humans, macaques, cats, short-tailed opossums, and mice). The timing of developmental transformations can be used to find corresponding ages across species. This figure is modified from Workman et al. (2013) as well as Charvet and Finlay, 2018. The event scale is a common ordering of developmental events across all studied species, which ranges from 0 to 1. Modeling the timing of these developmental transformations can serve to identify equivalent ages across species. By controlling for overall changes in developmental schedules, we can identify whether NEFH expression increases for longer than expected in humans compared with mice. Also represented on this graph are interaction terms for corticogenesis and retinogenesis, which represent deviations in the timing of developmental programs across select species. Events with blacked borders represent extensions in cortical or retinal neurogenesis with respect to their time of occurrence of other developmental transformations. Corresponding ages for marmosets and other species are in Charvet and Finlay, 2018.

**Figure 4.**
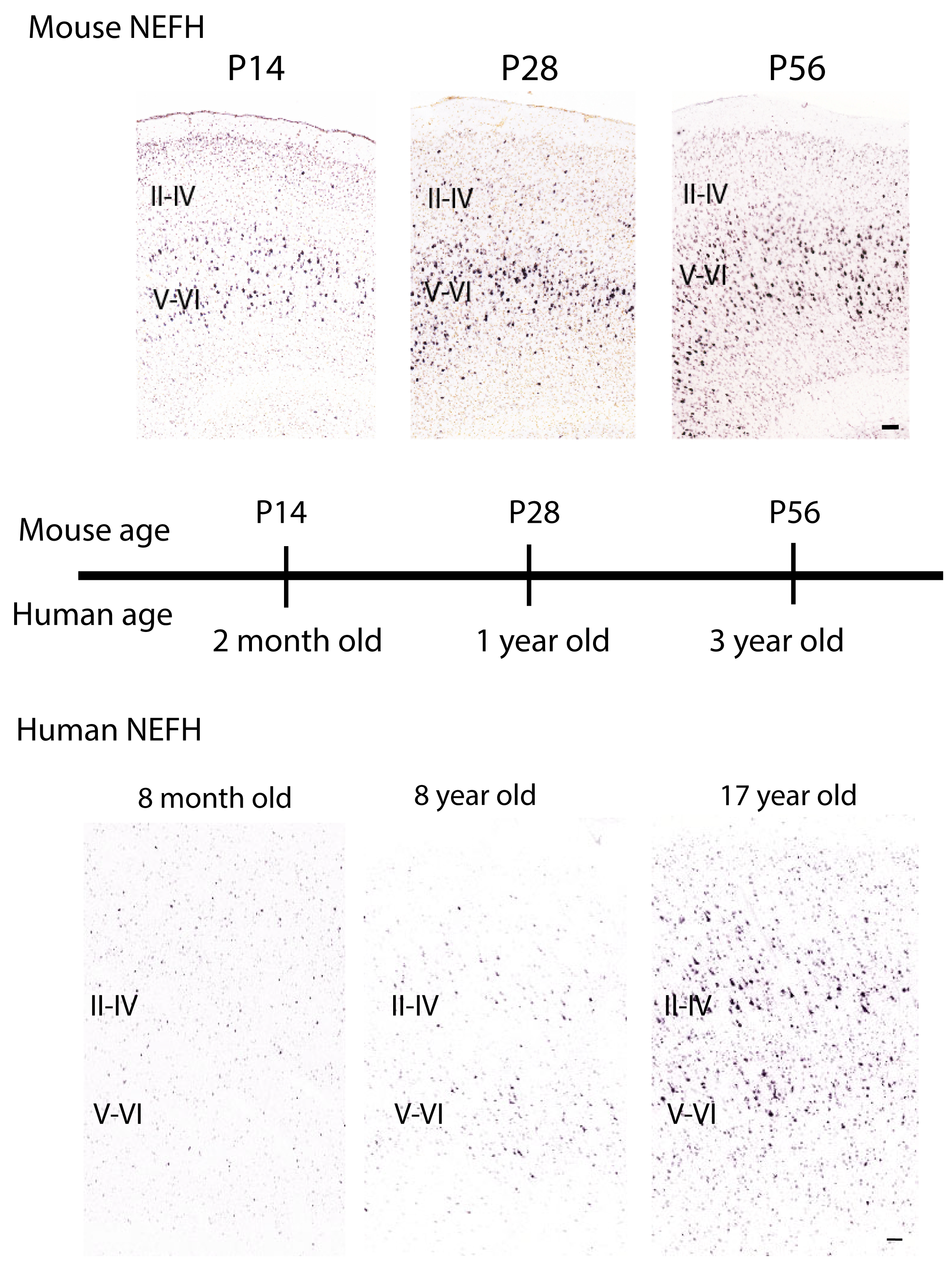
NEFH expression in the frontal cortex of mice and prefrontal cortical areas of humans at successive stages of postnatal development highlight that NEFH expression appears relatively stable fro P14 to P56 in mice. In contrast, NEFH expression is weak from 8 months to 8 years of age in human. NEFH expression in upper layers only clearly emerges by 17 years of age. Once overall differences in developmental schedules are controlled for, NEFH expression onset in upper layers appears protracted in humans relative to mice (see also **Fig. 5**). P: post-natal. Scale bars: 100 microns.

**Figure 5.**
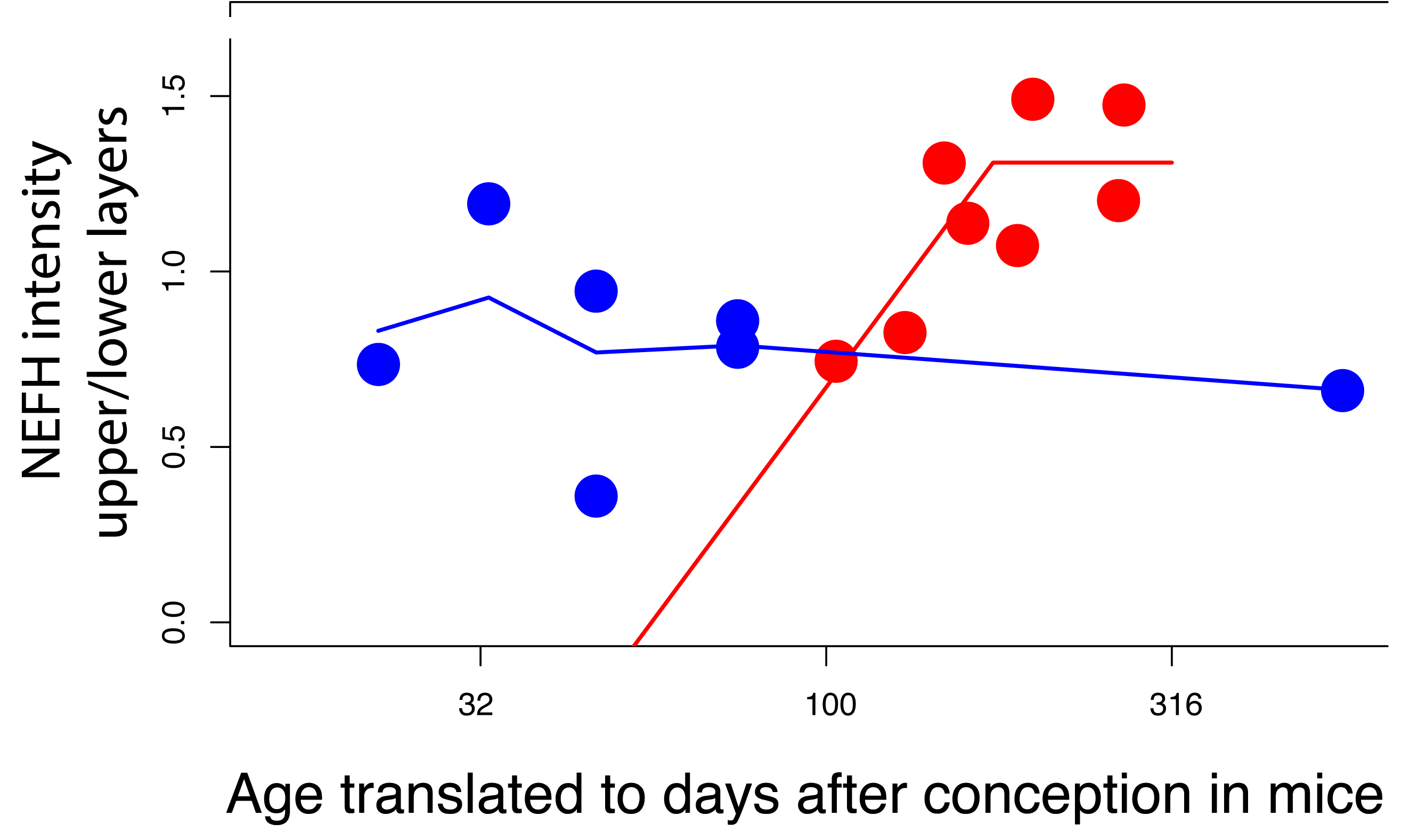
Developmental trajectories in the relative expression of NEFH expression in upper versus lower layers in the frontal cortical areas of humans and in mice, respectively. Ages of humans are translated to mice according to the translating time model (see **Fig. 2**). These data show that the relative expression of NEFH across layers steadily increases up to about 20 years of age in humans (∼ 5 to 6 month old mouse) and plateaus between 20 to 28 years of age. In other words, the expression of NEFH expression steadily increases in upper layers in humans but the expression of NEFH in upper versus lower layers remains relatively stable in mice. Testing for a linear plateau with easynls (R library packages) shows that the expression of NEFH in upper versus lower layers versus age is significant for a linear plateau (p<0.05; R^2^= 0.64) in humans. In mice, the expression of NEFH in upper versus lower layers remains relatively stable throughout the examined ages. The relative expression of NEFH does not reach a plateau in mice (p>0.05). A smooth spline fit through the data suggests that NEFH expression in upper versus lower layers may peak at around postnatal day14 in mice but the relative expression of NEFH remains relatively stable while the relative expression of NEFH steadily increases in humans. Taken together, these data highlight important heterochronic shifts in NEH expression across layers in both species. These data are consistent with those shown in **Fig. 1G**.

### Comparative analyses of NEFH expression across layers

A selective increase in NEFH expression in upper layers of the cortex would imply modifications to cortico-cortical pathways because layer II-III neurons preferentially form cortico-cortical pathways (Gilbert et al., 1975; Kennedy and Bullier, 1985; Barbas, 1986; Nudo and Masterton, 1990; Hof et al., 1995). To test the hypothesis that primates possess evolutionary changes in long-range cortico-cortical pathways relative to mice, we quantified differences in the relative expression of NEFH between upper (layers II-IV) versus lower layers (layers V-VI) in the primary visual cortex and frontal cortical areas of humans and mice. Age and samples used are listed in **Table 1**. We quantified the density of NEFH mRNA expression levels from sections in upper and lower layers of the primary visual cortex and frontal cortical areas of humans and mice. When comparing the relative intensity of NEFH expression in upper versus lower layers of adult brains (see **Fig. 1K, O**), we selected to compare NEFH expression densities in the primary visual cortex in humans between 8 to 41 years of age and mice between post-natal day 56 to 4 months of age. We also selected to compare NEFH expression densities in frontal cortical areas in humans between 17 to 57 years of age and in mice from post-natal day 28 to 24 months of age. As these ages, NEFH expression diverges in its pattern of expression between the two species (**Fig. 1 G**). We also compare temporal changes in NEFH expression between upper and lower layers across humans and mice (see **Fig. 5**).

We downloaded images of NEFH mRNA labeled sections made available by the Allen Brain Atlas. We placed a rectangular grid through sections that span the two cortical areas of interest (i.e., primary visual cortex, frontal cortical areas). We use this grid to randomly select sites along the cortical surface to measure the density in the relative expression levels of NEFH in upper and lower layers in the two species. Randomly-selected frames were aligned along the cortical surface. Frame widths were 500 and 1000 µm in mice and humans, respectively. The height of frames varied with the thickness of upper and lower layers. We used cyto-architecture from adjacent Nissl-stained section in humans, and in mice, as well as atlases to define cortical areas, as well as upper (i.e., layers II-IV) and lower layers (i.e., layer V-VI). Upper layers were distinguished from lower layers based on a small cell-dense layer IV evident from Nissl-stained sections. This approach is similar to that used previously to quantify upper and lower layer neuron numbers in primates and rodents (Charvet et al., 2015, 2017ac).

To compute the relative density of NEFH expression in upper versus lower layers in humans and in mice, we binarized images and computed the relative intensity of NEFH expression relative to the background in upper and lower layers, seperately. This approach permits quantifying the relative area occupied by NEFH expression in upper and lower layers, separately. We then computed the relative intensity of NEFH expression in upper versus lower layers in each individual (**Fig. 1G**). We, then, averaged NEFH relative intensity values for each specimen and tested for significant differences in NEFH expression between species with the use of a t test (**Table 1**). We used 4 to 12 frames per specimen and two to six sections per specimen to measure the relative densities in NEFH expression in upper and lower layers. To ensure the validity of our results, we used single cell RNA sequencing data-set extracted from the mouse primary visual cortex aged 6 to 8 weeks of age, as well as those obtained from the frontal cortex (see **Fig. 1, 6**). We computed the number of NEFH+ cells across layers of the mouse primary visual cortex or the frontal cortex (**Fig. 1L, O**). Data acquisition and cell type assignment of primary visual cortex neurons are described in detail and follow that of Hrvatin et al., 2018 (GSE102827). Data acquisition and cell type assignment of frontal cortex neurons are described in detail in Saunders et al., 2018 (GSE116470). Cell type assignments from Saunders et al., 2018 are shown in **Fig. 6**.

**Figure 6.**
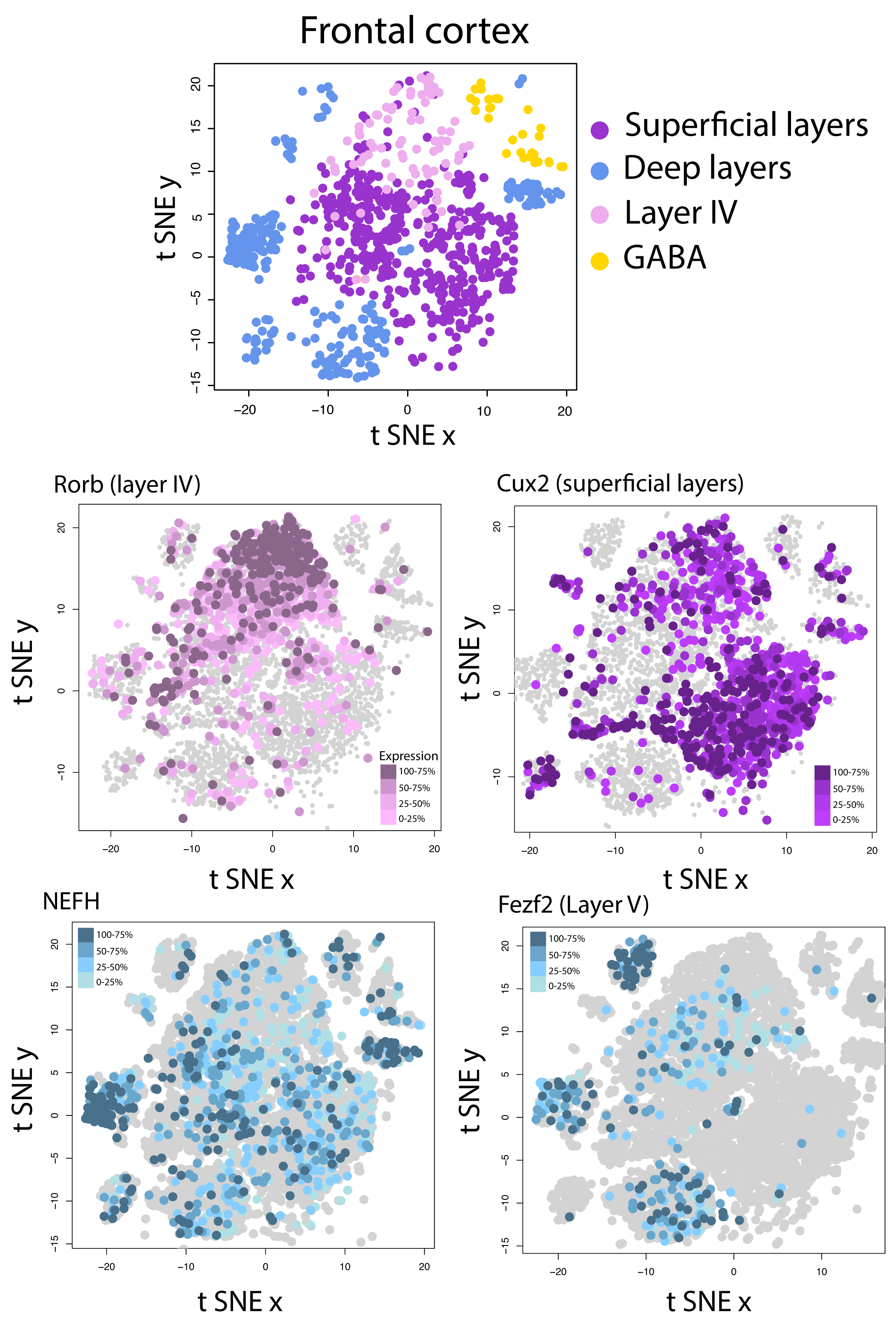
(A) t_SNE plots of RNA expression from single cells extracted from the frontal cortex of mice identifies clusters of cell populations. Rorb (B), Cux2 (C), and Fezf2 (D) are used as markers for layer IV, superficial (layers II-III), and lower layers (layer V), respectively. The expression of these marker genes coincides with the population identified in (A). NEFH expression appears to be well expressed in clusters identified as lower layers (e.g., Fezf2+ clusters), and less expression in clusters, which likely correspond to upper layers. To identify whether NEFH labeled neurons are preferentially observed in superficial versus lower layers, we compute the number of NEFH+ labeled cells in superficial and lower layers (see **Fig. 1P**). We here exclude glial cells from the analyses, and focus exclusively on neurons. Data and identified clusters were from generated by Saunders et al. 2018. Strength of gene expression is color-coded. Strongly expressed genes are darker. Shades of colors represent expression levels, which are binned according to quantiles.

## Results

### Overall fiber organization

We use high-resolution diffusion MR tractography to detect fiber pathways coursing across the brain of macaques, humans, marmosets, and mice. These tractographies show interwoven cortical white matter fibers coursing in different directions as assessed from the RGB code applied to the vector of fibers through the cortex of marmoset, macaques, human, and to a lesser extent in mice (**Fig. 1A**). Axial slice filters set though the cortex of these three species highlights a number of pathways coursing across the anterior to posterior and medial to lateral axis of the marmoset, macaque, and human cortex (**Fig. 1A-D**). The macaque cortex appears to have many more interwoven cortical pathways than the marmoset. A similar situation is observed in the human brain where a large number of pathways are observed coursing in a number of directions through the white matter of the cortex. In other words, the examined primate species possess a multitude of fibers according to the standard (red, green, blue) RGB code, applied to the vector to the direction of pathways. In mice, only the fiber (i.e., cingulate bundle) and pathways coursing in lateral inferior regions are observed coursing across the anterior to posterior of the isocortex (**Fig. 1D**). A few fibers are also observed coursing principally across the medial to lateral axes of the cortex. At least some, if not most, of these fibers likely comprise the corpus callosum (**Fig. 1D**). The observation that macaques, marmosets, and humans appear to possess many more muti-dimensional pathways coursing across the axes of the cortex compared with mice led us to test whether there are species differences in long-range connection patterns between some of these species.

NEFH is one of the few markers that can be used to detect neurons with thick and long-range axons (Hof et al., 1995). We compare NEFH expression levels between the frontal and visual cortex of primates and mice. In the primary visual cortex, the normalized log-based NEFH expression values are significantly higher in macaques (x= 2.85, SD= 0.12, ks test: p<0.05, n=6), chimpanzees (x= 2.93, SD=0.10, n=5, ks test: p<0.05), and humans (x=2.96, SD=0.12, n=5, ks test: p<0.05; **Fig. 1**) compared with mice (x= 2.93, SD=0.10, n=5, **Fig. 1E**). In the prefrontal cortex, the normalized log-based NEFH expression levels are significantly higher in macaques (x= 2.48, SD= 0.15; p<0.05, n=5) and humans (x= 2.77, SD= 0.13, n=6, p<0.05; **Fig. 1F**), but not in chimpanzees (x=2.76, SD=0. 29, n=6, ks test p=0.108), compared with mice (x=2.48, SD=0.15, n=5). The finding that NEFH expression values are greater in primates than in mice suggests major changes in long-range cortically projecting neurons between the two taxonomic groups.

To further validate the observed increases in NEFH expression between primates and mice, we investigate the developmental trajectories of NEFH expression levels in the frontal cortex of humans and mice. We rely on the timing of developmental transformations to find corresponding time points between model organisms and humans (Workman et al., 2013). Such an approach permits controlling for variation in developmental schedules across species in order to identify whether select developmental programs occur for an unusually long time in a given taxanomic group (Clancy et al., 2001; Charvet and Finlay, in press). To that end, we test whether a linear plateau captures NEFH expression over developmental time in the frontal cortex of both species. We then assess whether NEFH expression in the frontal cortex reaches a plateau significantly later in humans than in mice (**Fig. 1G**). NEFH expression in the frontal cortex increases with age to reach a plateau at 12 years of age in humans (easynls, model=3, adjusted R^2^=0.954, p<0.01; n=7); and reaches a plateau about 22 days after birth in mice (easynls, model=3, age of plateau=PCD 40.5; adjusted R^2^= 0.86; p<0.01; n=21).

After controlling for variation in developmental schedules between the two species, NEFH expression reaches a plateau significantly later than expected in humans. That is, NEFH expression steadily increases until 12 years of age in humans. NEFH expression reaches a plateau at post-conception day 40.5 (i.e., about 22 days after birth) in mice; **Fig. 1G**). According to the translating time model, a mouse on post-conception day 38.5-42.5 (i.e., postnatal day 20-24) is approximately equivalent to a human at 1 year of age (post-conception day 517; lower 95% CI: post-conception day 399; upper 95% CI: post-natal day 671; Workman et al., 2013; http://translatingtime.org). If temporal changes in NEFH expression were conserved between humans and mice, NEFH expression should reach a plateau at approximately 1 year of age. Yet, the plateau in NEFH expression in the frontal cortex of humans is achieved at 12 years of age in humans, which is 11 years later than expected, and well above the confidence intervals generated from the translating time model. The increase in NEFH expression follow a protracted developmental time course in humans relative to the timing of most other developmental transformations (**Fig. 1G**). The protracted increase in NEFH expression is concomitant with increased frontal cortex NEFH expression in humans relative to mice (**Fig. 1G**). At its plateau, NEFH expression in the frontal cortex is 1.6 times greater in humans (**Fig. 1G-H**), and NEFH mRNA levels in the frontal cortex become significantly greater in humans (x=1.83; SD=0.13; n=3) than in mice (x=0.14; SD= 0.05; n=11, p<0.05; ks.test; **Fig. 1H**). Thus, the protracted increase in NEFH expression is concomitant with an increase in NEFH expression in the adult human frontal cortex. These findings are consistent with the notion that there are major differences in cross-cortically connecting patterns between primates and mice, and these observations agree with those obtained from diffusion MR tractography, which together highlight major modifications to long-range cortically neurons between primates and mice. However, these data do not demonstrate whether the increase in NEFH expression observed in primates is due to an increase in NEFH expression in upper or lower layers

We next assess whether there is variation in NEFH expression across cortical layers in humans and in mice. A relative increase in the expression of NEFH in upper layers of the cortex in primates suggests an increase in the number of cortico-cortical projecting neurons because cortico-cortical pathways preferentially emerge from layer II-III neurons. Importantly, however, some lower layer neurons do form cortico-cortical pathways. A previous report showed that the expression of NEFH appears increased in upper layers of the primary visual cortex of humans relative to mice (Zeng et al., 2012; **Fig. 1**. I-J). To assess these differences quantitatively, we measured the area occupied by NEFH labeling in upper and lower layers in the primary visual cortex and the frontal cortex of humans and in mice. We then computed the density of NEFH expression in upper versus lower layers in both of these species in order to control for variation in background labeling across samples. A t test on the NEFH mRNA expression densities show that NEFH expression is relatively increased in upper versus lower layers in frontal cortical areas in humans (x=1.28; SD=0.17, n=6) compared with mice (x=0.68, SD=0.23, n=6, t= 5.11, p<0.05, **Fig. 1K**). Similarly, NEFH expression is significantly more expressed in the primary visual cortex of humans (x= 1.345, SD=0.40, n=5) than in mice (x=072, SD=0.22, n=4, t= 3.04, p<0.05; **Fig. 1O**).

We identify species differences in developmental trajectories in NEFH expression in frontal cortical areas in humans and in mice, with a protracted increase in NEFH expression in upper layers in humans relative to mice after controlling for overall changes in developmental schedules (**Fig. 3, 4**). NEFH expression steadily increases up and subsequently plateaus at 20-28 years of age (equivalent to about 5 months of age in mice). In contrast, expression of NEFH in upper versus lower layers remains relatively stable throughout examined ages in mice. Taken together, these data highlight important heterochronic shifts in NEH expression across layers between the two species.

To confrrm that NEFH expression is indeed relatively reduced in upper layers in mice, we use a previously published single cell RNA sequencing data-set extracted from the primary visual cortex. The authors used the t-Distributed Stochastic Neighbor Embedding approach (tSNE) to identify sub-populations of cell types within the primary visual cortex. Such an approach identified excitatory neurons across different layers of the mouse visual cortex. We use this cell type assignment to test whether NEFH expression is indeed poorly expressed in excitatory in upper layer neurons. Consistent with our analyses comparing NEFH expression across upper and lower layers, the vast majority of NEFH labeled excitatory neurons are indeed located in lower layers (i.e., layer V-VI) rather than in upper layers in mice (i.e., layers II-III; **Fig. 1L, O**). This situation holds for the primary visual cortex as well as the frontal cortex (**Fig. 1O**). These data suggest that primates may possess increased cortico-cortical pathways compared with mice.

### Cortico-cortical pathways in relation to cortical folding

To investigate the evolution of cortico-cortical pathways among non-human primates, we place ROIs through similarly-positioned cortical gyri in marmosets and in macaques (**Fig. 7, 8**). The placed ROIs across equivalent cortical regions (**Fig. 7**) identify a number of similarities but also some differences in connectivity patterns between the two species (**Figs. 8, 9**). Many of the highlighted fibers course across the anterior to posterior axis in macaques and marmosets, although some fibers course across the medial to lateral axes. Fibers coursing across the dorsal to ventral axes likely connect cortical and subcortical structures whereas fibers coursing across the anterior to posterior or medial to lateral axes likely form cortico-cortical pathways (Charvet et al., 2017a).

**Figure 7.**
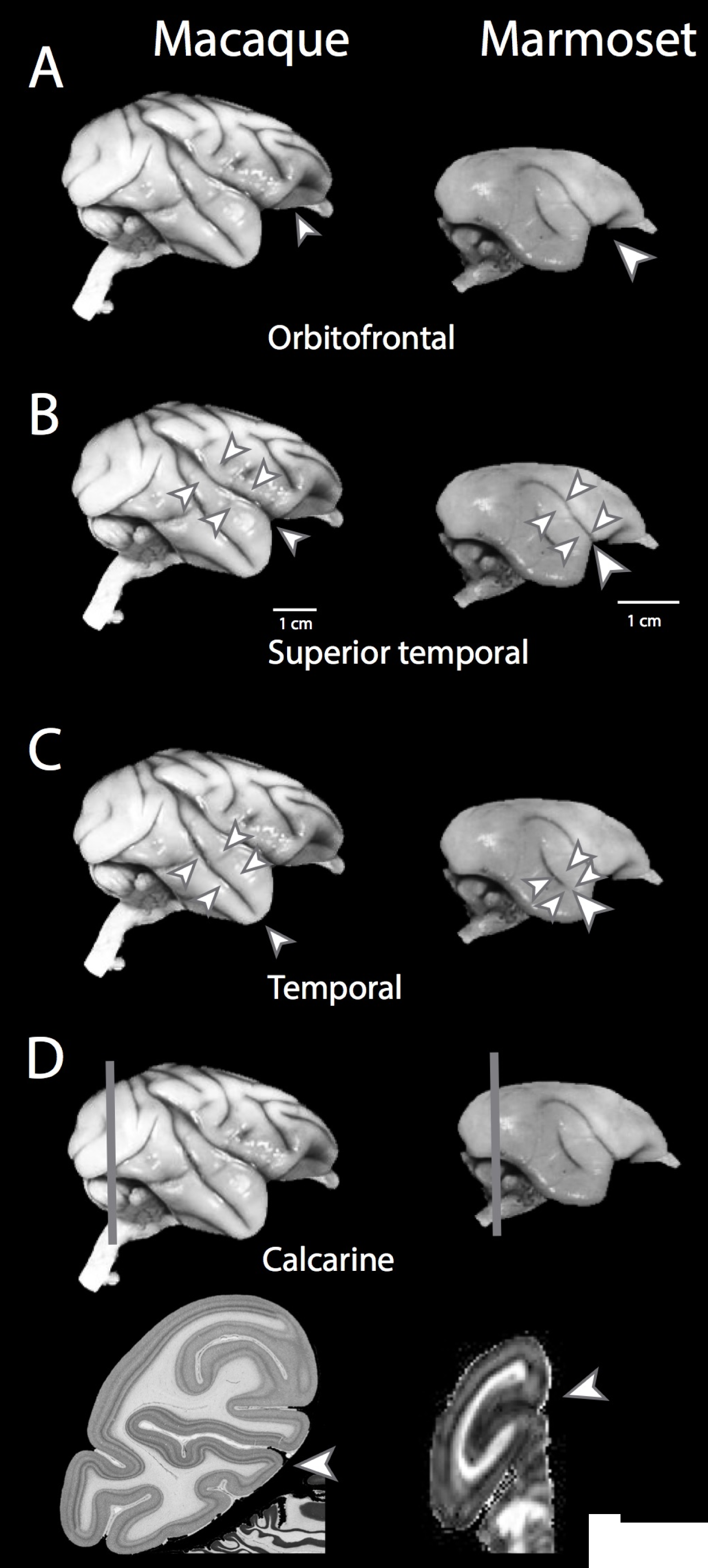
Arrowheads point to the location of ROIs placed through the macaque and marmoset cortex in order to identify cortico-cortical pathways coursing through these ROIs (see **Fig. 3, 4**). Arrowheads show the location of ROIs placed through the orbitofrontal (A), superior temporal (B) temporal (C) and calcarine sulci (D) of macaques and marmosets. (D) Vertical lines highlight approximate location through which coronal planes were taken. We use Nissl-stained sections through the macaque cortex and a coronal plane of a fractional anisotropy scan of the marmoset brain to highlight the location of ROIs through the calcarine sulcus.

**Figure 8.**
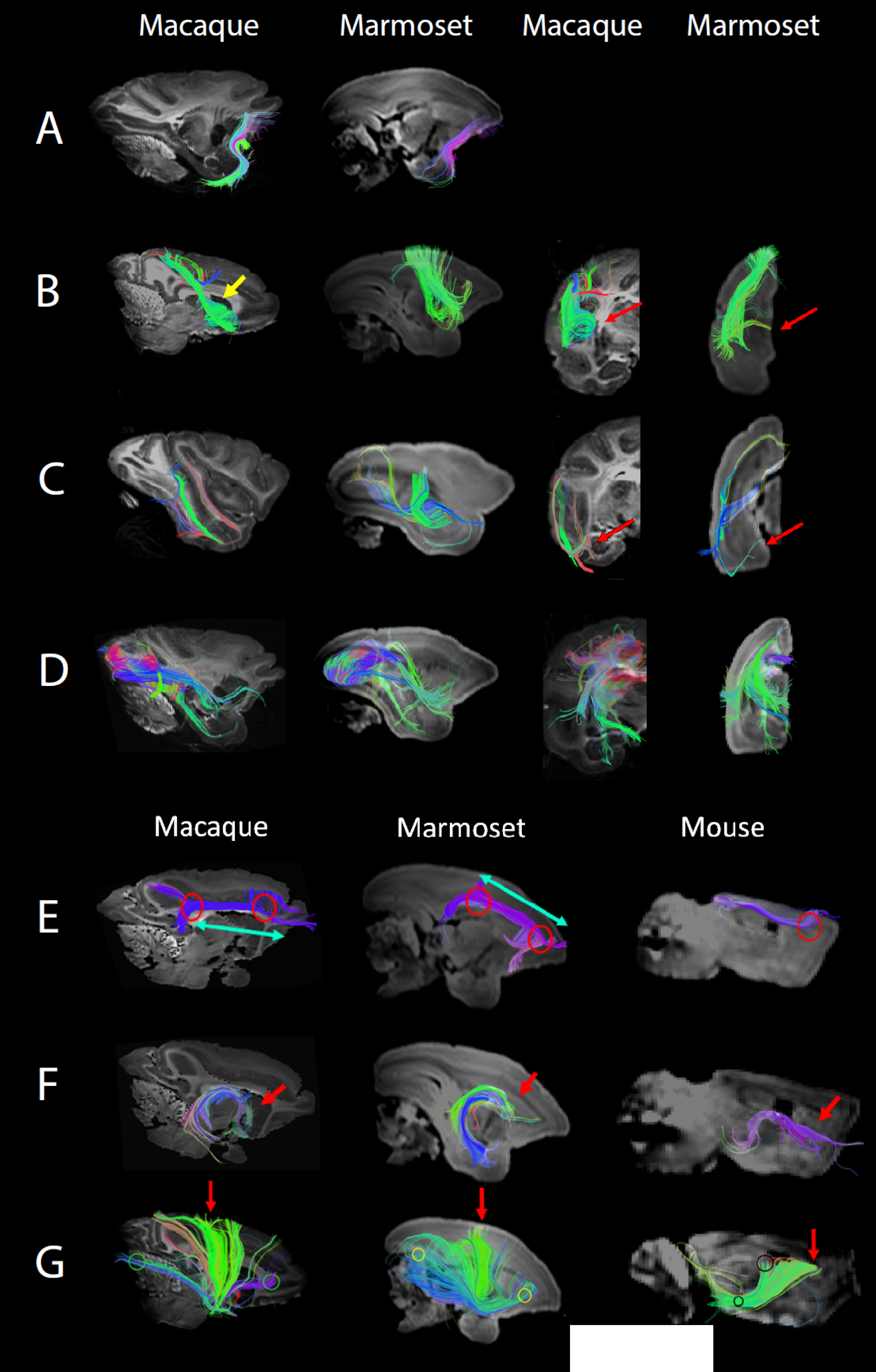
Panels (A-D) show fibers coursing through ROIs as shown in Figure 2. (A) ROIs show fibers coursing through the inferior orbitofrontal gyrus in macaques and marmosets. We define these fibers as the uncinate fasciculus given that they course between the frontal and temporal lobe. We note that the uncinate fasciculus is present in both macaques and marmosets and that these fibers principally course across the anterior to posterior axis (B) Sagittal and coronal slices of fibers coursing through the superior temporal areas. These fibers course across the medial to lateral axis and are observed in both macaques and marmosets. (C) Coronal and sagittal slices of fibers course through the temporal areas. In both macaques and marmosets, fibers course across the anterior to posterior axis between the frontal and temporal lobes. They satisfy our definition of the ILF. (D) Fibers show sagittal and coronal slices of fibers associated coursing through the calcarine sulcus. We successfully traced the cingulum bundle (E), fornix (F), and thalamocortical or corticothalamic (G) fibers in macaques, marmosets, and mice.

**Figure 9.**
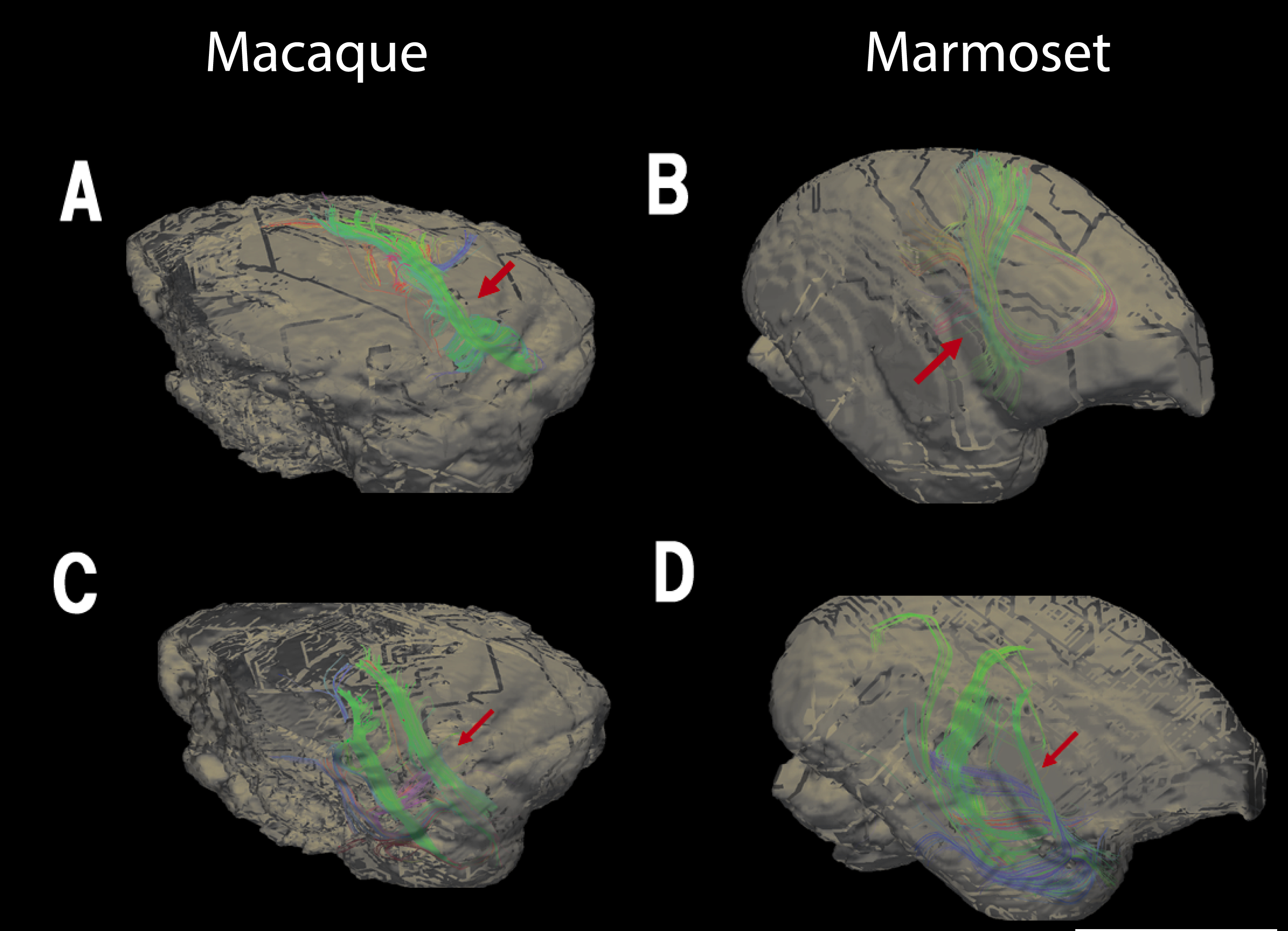
Panels (A) and (B) show the sagittal view of the fibers coursing tangentially to the superior temporal gryus. Panels (C) and (D) highlight sagittal views of fibers coursing tangentially to the temporal lobe gyrus. Panels (A) and (C) show the macaque brain. Panels (B) and (D) show the marmoset brain.

In both marmosets and macaques, ROIs placed in the orbitofrontal gyrus (**Fig. 7A**) show fibers coursing across the anterior to posterior axis between the frontal and temporal lobe, although the orbitofrontal gyrus in marmosets is shallow (**Fig. 8A**). Given that these fibers extend between the frontal and temporal lobes, we define these fibers as called the uncinate fasciculi. The uncinate fasciculus is observed in both macaques as in marmosets.

ROIs placed within the superior temporal gyrus (**Fig. 7B**) show that the fibers course tangentially to the main axes of sulci and gyri (see red arrows in **Fig. 9A, 9B**). These fibers course medially and perpendicular to the sulci (see red arrows in **Fig. 8B**). Macaques and marmosets vary slightly in their overall trajectories. The macaque fibers course between ventral regions of the frontal lobe (**Fig. 8B,** yellow arrow) and the parietal lobe whereas similar fiber pathways in marmosets course between the ventral to dorsal regions of the frontal lobe.

ROIs placed within the temporal gyrus (**Fig. 7C**) highlight fibers in macaques and marmosets coursing across the anterior to posterior axis in the temporal lobe, which extend dorsally towards the occipital lobe (**Fig. 3C**). We define these fibers as the inferior longitudinal fasciculus (ILF) because these white matter fibers course across the anterior to posterior axis within the temporal lobe and connect with the occipital lobe. We note the presence of the ILF is in marmosets and macaques (red arrows in **Fig. 9C, 9D**). ILF fibers extend fine branches coursing medially and perpendicular to the sulcus (red arrows in **Fig. 8C**). ROIs placed within the calcarine sulcus show similar fiber trajectories in both macaques and marmosets; in addition to the optic radiation, pathways coursing from/to the temporal and frontal lobes are identified (**Fig. 8D**). Taken together, these data demonstrate that macaques and marmosets possess a number of cortico-cortical pathways.

### Limbic and cortico-subcortical pathways

We clearly detect well-described pathways such as the cingulum, fornix, and thalamocortical and/or corticothalamic fibers (**Fig. 8E-G**). ROIs showed the cingulum, which courses across the anterior to posterior axis of the cortex across all three examined species. Marmoset and macaques possess branches coursing in two directions towards the frontal pole from the cingulum but the mouse cingulum fibers lack any type of branching extending across the anterior to posterior axis. Cingulum fibers in mice also appear thinner than in monkeys (see red circles in **Fig. 8E**). The macaque cingulum bundle spans its horizontal axis, while the marmoset cingulum bundle appears to be angled slightly upwards in its trajectory (see blue arrows in **Fig. 8E**). At the posterior end of the bundle, the macaque cingulum remains flat, while the marmoset bundle curves downward. In mice, the fornix lacked curvature in comparison to that of the monkey (see red arrows in **Fig. 8F**). The macaque and marmoset thalamocortical/corticothalamic fibers begin and terminate at similar corresponding locations (see red and yellow circles in **Fig. 8G**). In mice, fibers protrude towards the superior region of the frontal lobe (see black circles in **Fig. 8G**). In the primate brains, the fibers protrude in different directions from the origin point and span more cortical territory (see blue arrows in **Fig. 8G**). Thus, diffusion MR tractography successfully traces these pathways despite variation in brain geometry and size across the examined species.

We confirm our tractography results with those obtained from tract-tracers, made available by the Allen Brain Atlas (**Fig. 2;** Oh et al., 2014). The injection site in the Allen Brain Atlas ROIs from diffusion MR tractographies in the corresponding regions identify the cingulum bundle (**Fig. 2A, red arrows**), a part of the fornix (**Fig. 2B, red arrows**), a part of the thalamocortical pathways (**Fig. 2C, red arrows**), and a part of the corpus callosal pathways (**Fig. 2D, red arrows**). Lateral long-range pathways detected from diffusion MR tractography are concordant with those obtained from tract-tracers (**Fig. 2A, 2B, yellow arrows**).

## Discussion

In this study, we consider the three-dimensional perspective offered from high-resolution diffusion MR tractography to investigate the evolution of cortical pathways across euarchontoglires. Despite some species differences in branching patterns and variation in the number of cortical pathways coursing in different directions in macaques versus marmosets, our results highlight many similarities in patterns of connectivity between the two species (Zhang et al., 2013). At least some of these fibers we describe emerged early in primate evolution with or possibly before the emergence of haplorrhine primates (i.e., monkeys, apes). Our comparative analyses of NEFH expression coupled with diffusion MR tractography demonstrates that primates deviate from mice in possessing alterations in cortico-cortical pathways.

### Methodological approaches

The rapid acquisition of information from non-invasive scans coupled with the three-dimensional perspective offered from diffusion MR tractography provides a high-throughput approach with which to study the evolution and development of connections across species (Schwartz et al., 1991; Takahashi et al., 2010ab, 2012; Li et al., 2014; Song et al., 2015). Diffusion MR imaging, as with any other method, does have some limitations. For instance, diffusion MR tractography has difficulty resolving the precise terminations of pathways within the grey matter (Jones et al., 2013), which is especially challenging when comparing brains differing in size and structure. Diffusion MR tractography may also preferentially identify pathways based on certain characteristics (axon length, myelination) and neglect to identify other fibers (Reveley et al., 2015; Schilling et al., 2018). How best to resolve crossing fibers is an ongoing venue for research (Wedeen et al., 2008; Schilling et al., 2017; Maier-Hein et al., 2017). Diffusion spectrum and high angular resolution diffusion imaging can theoretically resolve crossing fibers (e.g. Takahashi et al., 2010, 2011, 2012), as well as pathways in poorly myelinated regions (Wilkinson et al., 2017). Indeed, our diffusion MR tractography successfully traced a number of well-characterized pathways. That is, we identified the fornix, cingulum bundle, thalamo-cortical or cortico-thalamic pathways with diffusion MRI tractography, which validates its use for cross-species comparisons. However, due to the potential limitations of diffusion MR tractography, we do not focus on detecting the precise terminations of pathways within the grey matter. Rather, we identify pathways coursing through similarly-positioned cortical gyri and sulci. It is also for these reasons that we adopt a descriptive rather than a quantitative approach, and we integrate observations from diffusion MR tractography with comparative analyses of NEFH expression.

### Evolution of long-range cortico-cortical pathways

The results from the present study quantitatively confirm that NEFH expression, a marker that identifies neurons projecting over long distances, is more strongly expressed in select cortical areas of primates compared with mice. We assessed whether the increase in NEFH expression in the primate cortex may result, at least in part, from an increase in NEFH expression in upper layers because layer II-III neurons preferentially form cortico-cortical pathways. Indeed, NEFH is significantly more expressed in upper layers of the cortex in humans than in mice. Our findings are in agreement with those of Zeng et al., 2012 who reported that NEFH is preferentially expressed in upper layers of the primary visual cortex in humans compared with mice. However, the study by Zeng et al., 2012 does not distinguish whether NEFH expression is increased in humans or whether the NEFH expression is shifted from lower to upper layers in humans. The present study demonstrates that NEFH expression is significantly increased in the frontal cortex and the primary visual cortex of primates. The increase in NEFH expression is concomitant with a relative increase in upper layer neurons in humans. These findings are consistent with the notion that humans possess increased cortico-cortical pathways compared with mice. Observations from diffusion MR data tractography highlight that the primate cortex contains a number of cortico-cortical pathways coursing in various directions through the white matter of the cortex. This is in contrast with mice where only a relatively few cortical pathways (e.g., corpus callosum, cingulum) are observed coursing across white matter of the isocortex (**Fig. 1**). Thus, the data from diffusion MR tractography and NEFH mRNA expression converge to show that primates possess increased cortico-cortical projecting neurons compared with mice.

### Evolution of long-range cortico-cortical pathways in primates

We set ROIs through the cortex of primates to identify long-range cortical pathways. The vast majority of these ROIs identified pathways forming cortico-cortical pathways rather than pathways connecting cortical and sub-cortical structures, which is consistent with the notion that primates possess a large number of cortico-cortical pathways (**Fig. 8**). We observed fibers coursing across the anterior to posterior direction of the cortex in macaques as in marmosets. Those include the ILF and the uncinate fasciculus. The ILF may consist of a series of fibers coursing across the anterior to posterior axes of the cortex and/or a series of long association fibers coursing between the occipital and temporal lobes (Tusa and Ungerleider 1985; Catani et al., 2003; Schmahmann et al., 2007). We also observe fibers coursing from the frontal to temporal lobes, which we define as the uncinate fasciculus. We here do not use diffusion MR tractography to detect the precise start or end location of these fibers. Previous tract-tracer studies and diffusion MR tractography studies noted that the ILF and uncinate fasciculus are present in humans and macaques but that it is either small or non-existent in rodents, or carnivores (Schmahmann et al., 2007; Schmahmann and Pandya, 2009; Takahashi et al. 2010ab, Charvet et al., 2017a). The finding that the ILF and uncinate fasciculi are present in marmosets suggests that the ILF and uncinate fasciculus evolved relatively early in primate evolution, with or before the emergence of haplorhine primates (i.e., monkeys, apes). It would be of interest to investigate whether the ILF and uncinate fasciculus are present in strepsirhine primates (e.g., lemurs) to trace the emergence of the ILF and uncinate fasciculus within the euarchontoglire lineage.

### Developmental source of variation in connectivity patterns

The observed differences in cortico-cortical pathways between primates and rodents can be traced back to developmental differences that serve to generate variation in the cortical composition between primates and rodents. Cortical neurogenesis occurs for an unusually long time in primates compared with rodents, even after controlling for overall variation in developmental schedules (Clancy et al., 2001; Charvet et al., 2017b). The consequence of extending the duration of cortical neurogenesis during development is that primates possess disproportionately more cortical neurons in adulthood. Most of the increase in cortical neuron numbers is accounted for by upper layer neurons (layers II-IV; Charvet et al., 2015, 2017a,b,c; Srinivasan et al., 2015, Atapour et al., 2018). Cortical neurogenesis onset is roughly similar between primates and rodents, but cortical neurogenesis is extended for longer than expected considering the timing of most other developmental transformations. As a result, layer IV and layer II-III neurons are produced for much longer than expected in primates than in rodents (Clancy et al., 2001; Charvet et al., 2011; Workman et al., 2013; Charvet et al., 2017b). Accordingly, extending the duration of cortical neurogenesis should lead to a concomitant amplification of local circuit layer IV neurons as well as layer II-III neurons forming long-range cortico-cortical pathways (Cahalane et al., 2014). Whether these two cell populations indeed co-vary across species has not been demonstrated quantitatively. Previous work comparing the relative contribution of local circuit layer IV neurons and long-range layer II-III cortico-cortical pathways across cortical areas highlight that these two cell populations appear to vary in concert in primates (Barbas et al., 2018). That is, NEFH labeled neurons numbers appear to increase progressively with laminar differentiation (Barbas and Garcia-Cabezas 2016). Interestingly, laminar structure predicts patterns of connectivity across the primate cortex with highly differentiated cytoarchitectural variation giving rise to increased layer II-III long-range cortico-cortical pathways (Barbas and Garcia-Cabezas 2016; Barbas et al., 2018). The expansion of upper layer neuron numbers in primates might be due to both increased local circuit layer IV neurons and increased long-range cortico-cortical pathways originating from layers II-III.

Many of cortico-cortical pathways described in primates (e.g., ILF, superior longitudinal fasciculus, uncinate fasciculus, arcuate fasciculus; Schmahmann et al., 2007; Schmahmann and Pandya, 2009) course across the anterior to posterior axes. Many of these pathways are not readily observed in other mammalian species (Takahashi et al., 2010a,b; Charvet et al., 2017a). Thus, primates may deviate from other mammals in possessing increased anterior to posterior cortico-cortical integration. The direction of these pathways align, at least to some extent, with variation in upper layer neuron numbers under a unit of cortical surface area, with upper layer neuron numbers varying from high values in the posterior cortex to low neuron numbers per unit of cortical surface area towards the frontal cortex (Charvet et al., 2015). Thus, long-range anterior to posterior cortico-cortical integration observed in primates aligns with the spatial variation in neuronal numbers under the cortical surface.

### Cortico-cortical projections and gyrification

A number of studies and hypotheses have considered how developmental processes may generate evolutionary changes in cortical composition, as well as gyrification across species (Goldman-Rakic and Rakic, 1984; Sun and Hevner, 2014; Striedter et al., 2015). An influential hypothesis posits that tension generated by short-range axons pull interconnected cortical regions together and cause gyrification (Van Essen, 1997), although a number of other explanations have been offered. The differential growth across the cortex, the migration of neurons along radial glia, and/or the differential expansion of upper versus lower layer neurons are a few proposed mechanisms generating variation in cortical folding between species (Richman et al., 1975; Van Essen, 1997; Toro and Burnod, 2005; Xu et al., 2010; Chen et al., 2013; Striedter et al., 2015; Fernández et al., 2016; Tallinen et al., 2016; Razavi et al., 2017). Modifications to neurogenetic programs are well-documented sources of variation across species, and experimental manipulations of cell proliferation during development impact cortical composition, as well as cortical folding (Dyer et al., 2009; Clowry et al., 2010; Charvet and Striedter, 2008, 2011; Charvet et al., 2017ab; Lewitus et al., 2013; McGowan et al., 2012, 2013; Nonaka-Kinoshita et al., 2013; Borrell and Götz, 2014; Otani et al., 2016; Tallinen et al., 2016; Toda et al., 2016; Matsumoto et al., 2017; Shinmyo et al., 2017; de Juan Romero and Borrell, 2017). We suggest that the differential composition of neurons across the depth of the cortex may cause major differences in white matter tension between species. Variation in cortical tension may, in turn, differentially impact cortical folding between mammalian orders (Pillay et al., 2007; Zilles et al., 2013). Modifications to cortical composition and projection patterns evolve in concert, and may together lead to major differences in the extent of cortical folding between primates and rodents. Although the precise mechanisms generating buckling of the cortex is an ongoing matter of debate, it is likely that a number of coordinated factors, rather than a single factor, dictate the extent and spatial variation of cortical folds.

## Acknowledgments

This work was supported the Eunice Shriver Kennedy National Institute of Child Health and Development (NICHD) (R01HD078561, R21HD069001) (E.T.), the National Institute of Mental Health (NIMH) (R21MH118739) (E.T.), the National Institute of Neurological Disorders and Stroke (R03NS091587) (E.T.), as well as grant number 5P20GM103653 from the National Institute of General Medicine Sciences. This research was carried out in part at the Athinoula A. Martinos Center for Biomedical Imaging at the Massachusetts General Hospital, using resources provided by the Center for Functional Neuroimaging Technologies, NIH P41RR14075, a P41 Regional Resource supported by the Biomedical Technology Program of the National Center for Research Resources (NCRR), National Institutes of Health. This work also involved the use of instrumentation supported by the NCRR Shared Instrumentation Grant Program (S10RR023401, S10RR019307, and S10RR023043), and High-End Instrumentation Grant Program (S10RR016811). We thank Drs. Guangping Dai and Lana Vasung for providing technical assistance. Photos of mouse, macaque, and marmoset brains are from the comparative mammalian brain collection (http://neurosciencelibrary.org). Images of tractography from tract-tracers, NEFH in situ hybridization data from mice and humans were taken from the Allen Institute Website and the Brainspan atlas of the developing human brain. These data are available at http://developingmouse.brain-map.org and at http://www.brainspan.org, which are supported by the NIH Contract HHSN-271-2008-00047-C to the Allen Institute for Brain Science. The opinions in this article are not necessarily those of the NIH.

## Figure Legends

Table 1. List of ages and samples used for comparative analyses of NEFH expression, as well as diffusion MR tractography.

